# Cross-pathway integration of cAMP signals through cGMP and calcium-regulated phosphodiesterases in mouse striatal cholinergic interneurons

**DOI:** 10.1101/2023.10.07.560741

**Authors:** Ségolène Bompierre, Yelyzaveta Byelyayeva, Elia Mota, Marion Lefevre, Anna Pumo, Jan Kehler, Liliana R.V. Castro, Pierre Vincent

**Affiliations:** Sorbonne Université, CNRS, Biological Adaptation and Ageing, F-75005 Paris, France; H. Lundbeck A/S, Ottiliavej 9, DK-2500 Valby, Denmark; IGF, Univ. Montpellier, CNRS, INSERM, Montpellier, France

**Keywords:** Cholinergic interneurons, Striatum, cAMP, cGMP, calcium, Phosphodiesterases, Biosensor imaging

## Abstract

Acetylcholine plays a key role in striatal function, yet the intricate dynamics of cyclic nucleotide signaling which govern the firing properties of cholinergic interneurons (ChINs) have remained elusive. Since phosphodiesterases determine the dynamics of cyclic nucleotides, in this study, we used FRET biosensors and pharmacological compounds to examine phosphodiesterase activity in ChINs in mouse brain slices. Intriguingly, these neurons displayed strikingly low levels and slow cAMP responsiveness compared to medium-sized spiny neurons (MSNs). Our experiments revealed that PDE1, PDE3 and PDE4 are important regulators of cAMP level in ChINs. Notably, the induction of cGMP production by nitric oxide (NO) donors increases cAMP by inhibiting PDE3 - a mechanism hitherto unexplored in neuronal context. Furthermore, the activation of NMDA or metabotropic glutamate receptors increases intracellular calcium, consequently activating PDE1 and thereby decreasing both cAMP and cGMP. This interplay of phosphodiesterases enables the control of cAMP by the neuromodulatory influences of glutamate and NO. Remarkably, the NO/cGMP signal results in different effects: NO enhances cAMP in ChINs by inhibiting PDE3, whereas it reduces cAMP levels in MSNs by activating PDE2A. These findings underscore the specificity of intracellular signaling in ChINs compared to MSNs and show how the NO-cGMP pathway affects these various neuronal types differently. These observations have significant implications for understanding the regulation of the striatal network and the integration of dopaminergic signals and suggest innovative therapeutic strategies for addressing basal ganglia disorders with unmet medical need.

## Introduction

Basal ganglia play an important role in action selection, motor learning and reward, and striatal dysfunction is involved in various diseases such as Parkinson’s, Huntington’s, addiction. The vast majority of striatal neurons are GABAergic projection neurons called medium-sized spiny neurons (MSN), while about 1% consists in cholinergic interneurons (ChIN). Acetylcholine works in concert with dopamine to ensure the temporal coordination of striatal circuitry and therefore define a temporal windows for learning (1–3). By synchronizing MSN activity, ChIN are involved in changes and termination of movements (4–6) and behavioral learning (2).

Regular spontaneous firing of ChINs maintains a cholinergic tone throughout the striatum, and this tonic firing is interspersed with burst - pause - rebound sequences coincident with phasic dopamine release (7). Electrophysiological studies revealed that ChIN activity is controlled by various neuromodulators involving the cAMP signaling pathway such as noradrenaline (via β-adrenergic receptors) or dopamine (via D5 receptors) (8, 9). ChINs show large I*_h_* current, mainly mediated by the HCN2 channel (10, 11) which is under direct control of intracellular cAMP (12–14). The binding of cAMP to HCN channels shifts their activation curve, facilitating depolarization and increasing the firing frequency. In spite of its considerable importance, the control of cAMP signaling pathway in ChIN remains poorly understood.

cAMP is a highly dynamic intracellular second messenger, whose concentration varies rapidly and is determined by production by adenylyl cyclases and degradation by 3’,5’-cyclic nucleotide phosphodiesterases. There are numerous isoforms of phosphodiesterases that vary, for example, in enzymatic specificity (cAMP and/or cGMP), Km, subcellular targeting.(15–19) Moreover, their activity can be controlled by other second messengers, making these enzymes a key hub for cross-pathway regulations (20). Each cell type expresses a precise subset of phosphodiesterases leading to distinct integrative properties.

Our objective was to determine which phosphodiesterases are functionally present in ChINs and how their expression pattern might determine the integration of the cAMP signal. PDE1A is expressed in ChINs, as shown by immunocytochemical and single-cell transcriptomic analysis (22, 28). The PDE1 family degrades both cAMP and cGMP and is active only in the presence of calcium-calmodulin (29), which suggests a potential link between calcium signals and the regulation of intracellular cyclic nucleotide levels. PDE3A mRNA is expressed in ChINs, as shown by single-cell RNA sequencing and by the punctate labeling in in-situ hybridization (21, 22). PDE3 has a sub-micromolar Km for cAMP and is inhibited by cGMP (23–27). The widely expressed PDE4 family is also present in ChINs (22). PDE4 does not degrade cGMP. While PDE2A and PDE10A are highly expressed in the striatum, they are below detection level in ChINs (22, 33, 34). Whether the predicted phosphodiesterases actually contribute to the regulation of cAMP in ChINs is unknown.

Genetically encoded FRET biosensors, which provide a direct measurement of intracellular signals in a physiologically relevant context, have revealed the importance of phosphodiesterases in the dynamic regulation of cyclic nucleotides. We showed in MSNs in striatal mouse brain slices that PDE1B and PDE2A mediate the action of calcium or cGMP signals on cAMP (35, 36). The aim of this study was to use a similar approach to monitor the functional contribution of the various phosphodiesterases in the regulation of cAMP levels in ChINs, and to explore how these phosphodiesterases might convey signals from NO and glutamate to the cAMP signaling pathway.

## Methods

### Brain slice preparations

Mice (male and female C57BL/6J; Janvier labs) were handled in accordance with the Sorbonne University animal care committee regulations. Brain slices were prepared from mice aged from 7 to 12 days. Mice were decapitated and the brain was quickly removed and immersed in an ice-cold solution of the following composition: 125 mM NaCl, 0.4 mM CaCl_2_, 1 mM MgCl_2_, 1.25 mM NaH_2_PO_4_, 26 mM NaHCO_3_, 20 mM glucose, 2.5 mM KCl, 5 mM sodium pyruvate and 1 mM kynurenic acid, saturated with 5% CO_2_ and 95% O_2_. Coronal striatal brain slices of 300 µm thickness were cut with a VT1200S microtome (Leica). The slices were incubated in this solution for 30 minutes and then placed on a Millicell-CM membrane (Millipore) in culture medium (50% Minimum Essential Medium, 50% Hanks’ Balanced Salt Solution, 5.5 g/L glucose, penicillin-streptomycin, Invitrogen). The cAMP biosensor Epac-S^H150^ (36), cGMP biosensor cyGNAL (35) and calcium biosensor Twitch-2B (37) were expressed using the Sindbis virus as vector (38): the viral vector was added on the brain slices (∼5 x 105 particles per slice), and the infected slices were incubated overnight at 35°C under an atmosphere containing 5% CO_2_. Before the experiment, slices were incubated for 30 min in the recording solution (125 mM NaCl, 2 mM CaCl_2_, 1 mM MgCl_2_, 1.25 mM NaH_2_PO_4_, 26 mM NaHCO_3_, 20 mM glucose, 2.5 mM KCl and 5 mM sodium pyruvate saturated with 5% CO_2_ and 95% O_2_). Recordings were performed with a continuous perfusion of the same solution at 32°C.

### Chemicals and drugs

Solutions were prepared from powders purchased from Sigma-Aldrich (St Quentin-Fallavier, Isère, France). Tetrodotoxin (TTX) was from Latoxan (Valence, France). Other compounds and drugs were from Tocris Bio-Techne (Lille, France).

**Table 1:**
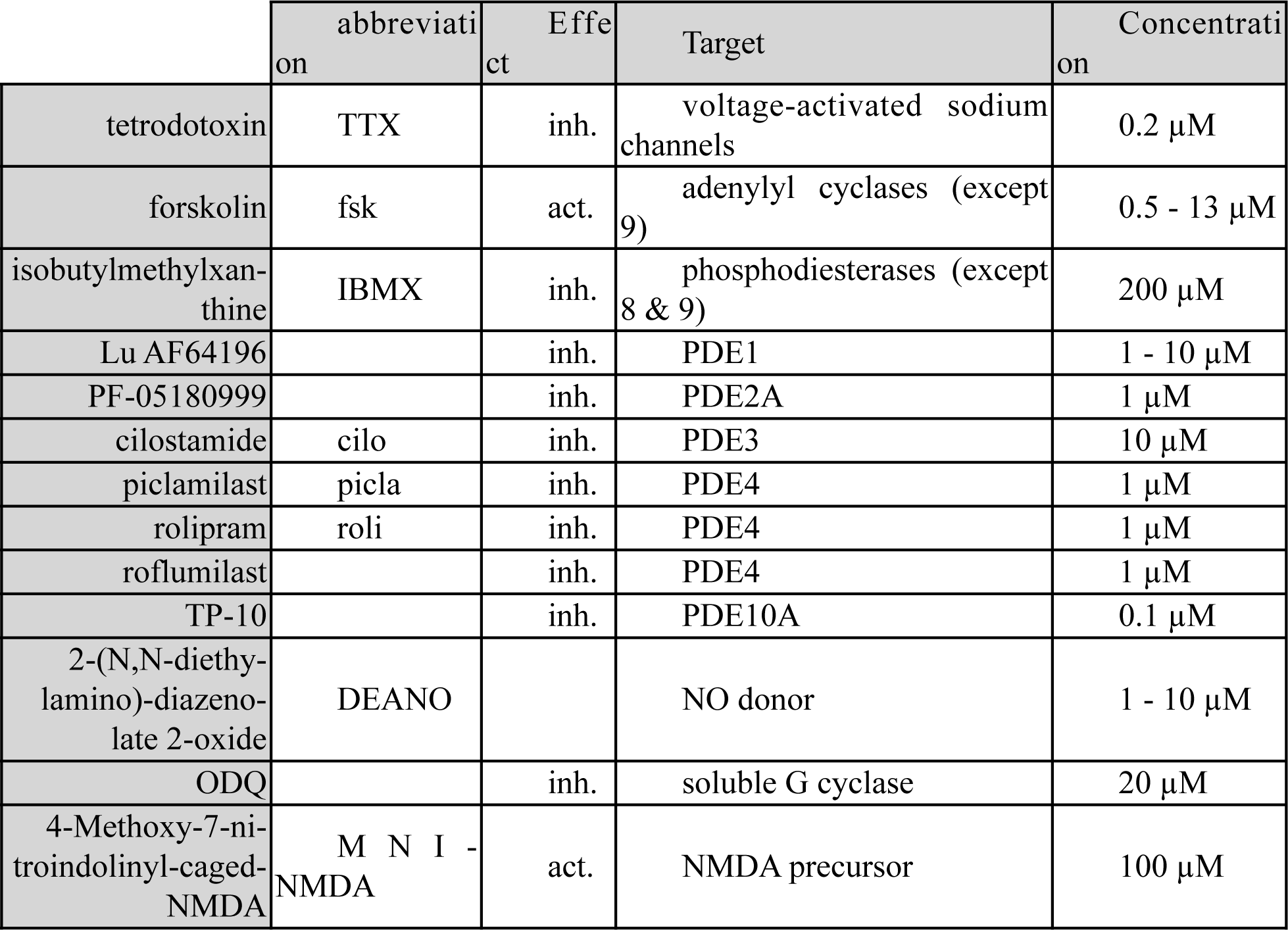
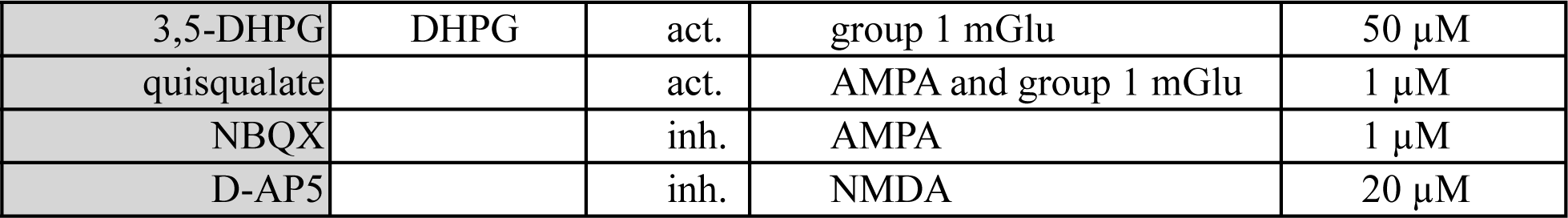
Drugs. List of the drugs used in this study. Effect indicates inhibitor (inh.) or activator (act.).

### Biosensor imaging

Imaging was performed in the dorso-medial part of the striatum. Wide-field images were obtained with an Olympus BX50WI or BX51WI upright microscope with a 40x 0.8 numerical aperture water-immersion objective and an ORCA-AG camera (Hamamatsu). Images were acquired with iVision (Biovision, Exton, PA, USA). The excitation and dichroic filters were D436/20 and 455dcxt. Signals were acquired by alternating the emission filters, HQ480/40 for donor emission, and D535/40 for acceptor emission, with a filter wheel (Sutter Instruments, Novato, CA, USA). The frequency of data acquisition was usually 1 image pair every 30 or 60 seconds, but was increased to resolve transient responses. To increase the number of recorded neurons, up to 3 focal depths were recorded for each time-point. Excitation power at the exit of the microscope objective was 400 to 1400 µW. Photo-release of NMDA from “caged” precursor MNI-NMDA was performed using a high power 360 nm light-emitting diodes mounted on the epifluorescence port of the microscope, for 0.5 s duration with 14 mW power at the exit of the microscope objective. Filters were obtained from Chroma Technology and Semrock. LED light sources (420 nm with a 436 nm excitation filter and 360 nm) were purchased from Mightex (Toronto, Canada).

Wide-field imaging allowed for the separation of ChINs signal from MSNs, provided that the infection level was kept low and no fluorescence overlap between neighboring neurons was observed. In a few cases, cells were excluded when basal ratio was elevated, when the response to maximal stimulation was lacking or when the neuronal morphology was altered (uneven cell contours). At the end of every biosensor imaging experiment, an image stack was acquired to ascertain the morphology of the recorded cells in 3 dimensions.

An application of TTX (200 nM) for 5 min preceded drug applications to stop any ongoing spontaneous activity; in some cases, this induced a decrease in Twitch-2B ratio, consistent with the ChIN exhibiting a tonic firing activity before TTX.

### Image analysis

Images were analyzed with a custom software, Ratioscope (39), written in the IGOR Pro environment (Wavemetrics, Lake Oswego, OR, USA) as described previously (40). No correction for bleed-through or direct excitation of the acceptor was applied. Biosensor activation level was quantified by ratiometric imaging: donor fluorescence divided by acceptor fluorescence for Epac-S^H150^ and cyGNAL, and acceptor divided by donor for Twitch-2B. The emission ratio was calculated for each pixel and displayed in pseudo-color images, with the ratio value coded in hue and the fluorescence of the preparation coded in intensity.

Biosensor concentration was estimated from the fluorescence intensity measured as follows. A vertical image stack was acquired at the end of each experiment. For each recorded neuron, the plane in the stack where the neuron showed the best contrast was selected. Since the whole region of interest encompasses some background or dimmer regions of the cell, the intensity was evaluated over the 20% brightest pixels within the region of interest in this plane of the stack. This value is corrected for camera offset, exposure duration, excitation power and shading correction, and expressed as counts per pixel per milliwatt of power excitation per second (c/ mW/pixel/s).

### Statistical analysis

Statistical tests were performed with the same Ratioscope package using built-in Igor Pro functions. Brain slices were considered independent. One experiment is defined as one brain slice being used in a protocol such as that illustrated in Figure 2A. Only a single experiment was performed on a brain slice. While most experiments were done with a single ChIN, if more than one was present (up to 4), their responses were averaged. Statistics were calculated per experiment (N); (n) indicates the total number of neurons per condition; (A) indicates the number of animals used for preparing all the brain slices used per condition. No outlier was excluded in this study. The alpha level for statistical significance was 0.05, indicated by * in figures and tables; n.s. indicates that the test showed no significant difference. For some conditions, the Shapiro-Wilk test rejected the hypothesis of normal data distribution and therefore non-parametric statistics were used for comparing ratio levels throughout this study. The Wilcoxon signed rank test was used to compare two measurements of paired data. The Friedman’s rank test was used to determine an effect when more than 2 conditions were compared. The Wilcoxon-Mann-Whitney test was used to test whether the treatment increased the ratio value for unpaired data.

### Estimates of biosensor activation level

Ratiometric imaging can be used to determine absolute analyte concentrations, with ratio values following a Hill equation from Rmin, the ratio in the absence of ligand, to Rmax, the ratio in the presence of saturating ligand (41). For Epac-S^H150^, considering the low cAMP tone in ChIN (see below), the basal ratio was assumed to be Rmin. Rmax was determined for each neuron by the final application of 13 µM forskolin and 200 µM IBMX, which is commonly accepted as a condition in which the biosensor is saturated with cAMP (42). An axis on the right presents cAMP concentration estimates based on these Rmin and Rmax value, a Hill equation with a Kd of 10 µM and a Hill coefficient of 0.77 (36, 43). With the cyGNAL biosensor, baseline ratio was considered equal to Rmin (see below); Rmax was determined for each neuron by the final application of 10 µM DEANO and 200 µM IBMX. An axis on the right presents cGMP concentration estimates based on these Rmin and Rmax value, a Hill equation with a Kd of 0.465 µM and a Hill coefficient of 0.8 (35). For calcium imaging, the Rmax value was difficult to determine since common manipulations aiming at maximizing intracellular calcium, such as NMDA, ionomycin or KCl applications, triggered ratio increases of inconsistent and unstable value. Calcium measurements are therefore presented as raw ratio values.

### Immunohistochemistry

After biosensor imaging, brain slices were fixed by overnight immersion in a 4% paraformaldehyde (PFA) in phosphate buffer (0.1 M, 4°C, pH 7.4). They were then stored at −20°C in a phosphate buffer containing 30% (v/v) glycerol and 30% (v/v) ethylene glycol, until they were processed for immunofluorescence. Choline acetyltransferase (ChAT) expression was detected with a goat polyclonal antibody (1:500; 4°C overnight, AB144p, Merck) and revealed with the secondary antibody Alexa Fluor 647 donkey anti-goat IgG (1:500, Jackson Immunoresearch) imaged with F39-628 filter for excitation, F37-692 filter for emission and F38-661 dichroic mirror (Olympus). Following the incubation of the primary antibodies (4°C, overnight incubation), brain sections were rinsed and incubated with the secondary antibody for 2 h at room temperature. Floating slices were rinsed and images were acquired with a wide-field microscope.

## Results

### Identification of Cholinergic Interneurons in a brain slice

We used genetically encoded biosensors to explore intracellular signaling events in individual neurons in the context of brain slices from mouse pups (Figure 1A). Sparse biosensor expression in neurons was obtained after overnight incubation of brain slices with the Sindbis viral vector as described previously (40). In the course of our previous studies on cAMP dynamics in MSNs, we were intrigued by the odd behavior of a small fraction of neurons in the imaging field which showed small and slow cAMP responses to forskolin (fsk, 13 µM, Figure 1B, left). In these neurons, the amplitude of the response to forskolin was 0.25 (N=47, n=69, A=21) of the maximal response while it reached 0.92 (N=38, n=105, A=21) in MSNs (Wilcoxon test, P<0.001). We verified that the moderate size of the cAMP response to forskolin did not result from a buffering effect of excessive biosensor expression. The biosensor expression level was estimated for each neuron from the fluorescence intensity of the cell body (see Methods) and revealed a fluorescence intensity in these neurons of 0.29 counts/mW/pixel/s (n=60), somewhat larger (t test, *P*<10^−6^) than in MSNs (0.18 counts/mW/pixel/s, n=87). However, there was no correlation between the amplitude of the cAMP response in these neurons and the biosensor expression level (Figure 1B, middle; n=60, r=-0.04), ruling out an artifactual effect of biosensor level on these neurons’ responsiveness to forskolin. The response to forskolin also appeared much slower in these neurons than in MSNs (Figure 1B, right, Wilcoxon test, P<0.001). The onset slope did not appear related to biosensor concentration (n=60, r=-0.17), again arguing against a buffering effect of biosensor on cAMP signal. These neurons’ morphology was also quite distinct from MSNs, with a cell body diameter of over 13 µM, an elongated cell body, an ovoid nucleus and wide proximal dendrites, suggesting they were cholinergic interneurons (44, 45). This morphological identification was confirmed by immunohistochemical labeling of Choline Acetyltransferase (ChAT) after biosensor imaging experiments (Figure 1C). 11 brain slices were fixed after the biosensor experiment and later processed for ChAT immunoreactivity. In these slices, 15 neurons were visually identified as ChINs during the biosensor recording session. All of these neurons showed a positive ChAT labelling.

**Figure 1:**
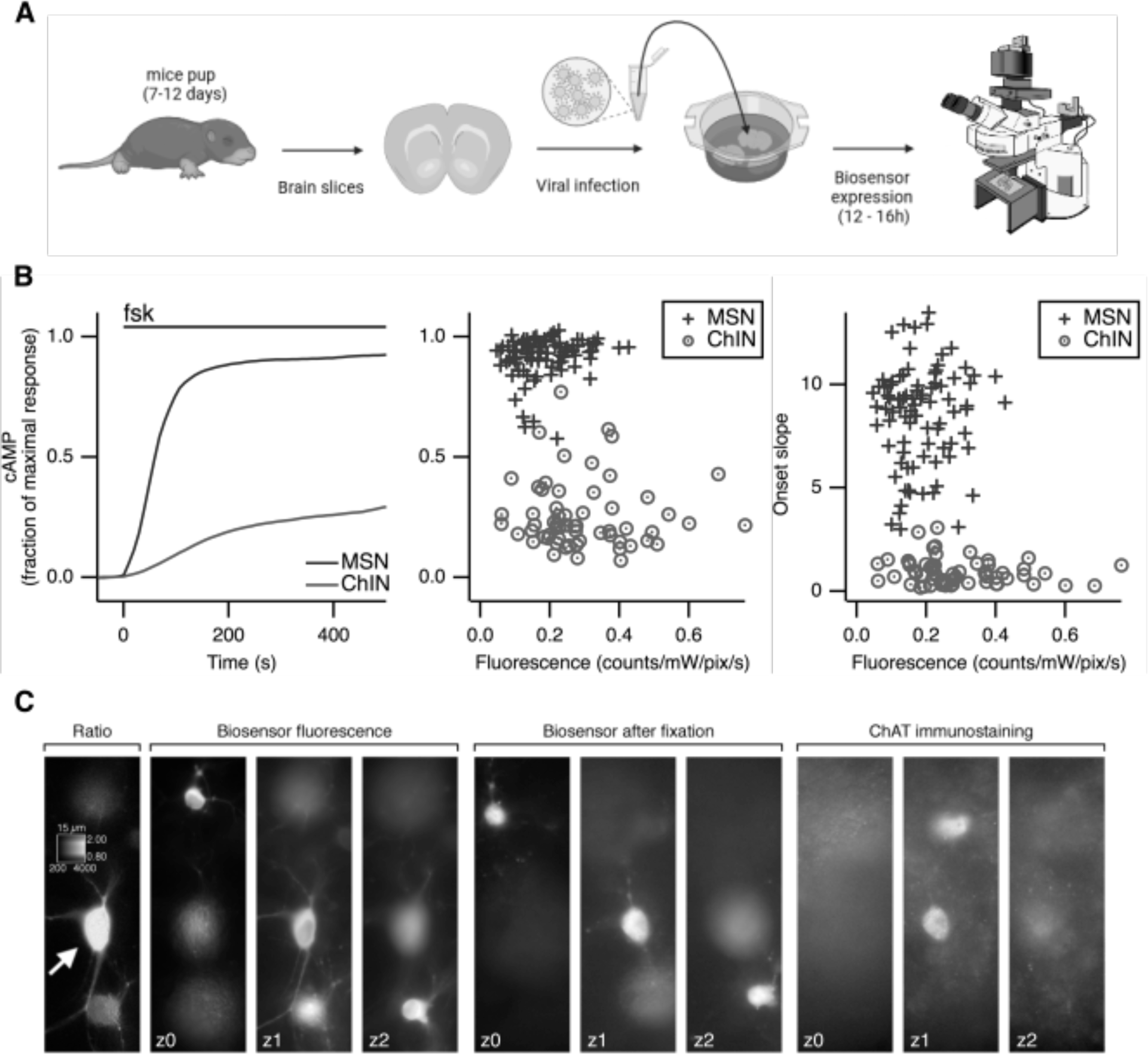
Biosensor imaging in mouse brain slices and identification of striatal cholinergic interneurons (ChIN) (A) Brain slices were prepared from 7-12 days old mice pups, and kept on a membrane over culture medium. The sindbis viral vector for biosensor expression was deposited over the slice. Fluorescence imaging was performed after overnight incubation. (B) cAMP responses to forskolin in striatal ChINs and MSNs, recorded with the Epac-S^H150^ biosensor for cAMP. Left: the responses to forskolin (fsk, 13 µM) were averaged per neuronal type; maximal response corresponds to the maximal cAMP level in response to fsk and IBMX (200 µM). Amplitude (middle graph) and onset slope (right graph) of the cAMP response to fsk, plotted as a function of biosensor fluorescence. (C) ChIN identification in brain slices. This brain slice was transduced for the expression of the Twitch-2B calcium biosensor. The pseudocolor image (left) displays the biosensor emission ratio in two MSNs and one ChIN (arrow). At the end of the recording, the slice was fixed and immunostained for Choline Acetyltransferase (ChAT). 3 different focal planes (z0, z1 and z2) are represented in the images in gray levels. From left to right: biosensor fluorescence during the biosensor imaging session, after slice fixation with PFA and after ChAT immunostaining. The arrow indicates a ChIN; the other neurons are putative MSNs except for one additional ChIN revealed by ChAT that did not express the biosensor (z1 image in ChAT immunostaining panel).

### PDE3 and PDE4 control cAMP levels in ChIN at rest

Considering the attenuated cAMP responses in ChINs, our first objective was to identify which phosphodiesterases are involved in the negative control of cAMP level. We performed wide-field microscopy, sometimes at several focal planes, to record more ChINs in the same field (Figure 2A, ChIN 0 and ChIN 1 on the first focal plane (z0 image) and ChIN 2 on the second focal plane (z1 image)). We first stimulated cAMP production with forskolin, which increased cAMP level moderately in ChINs, and then added the selective PDE3 inhibitor cilostamide (cilo, 10 µM) which increased the ratio in ChINs to a higher steady-state level. Further addition of the selective PDE4 inhibitor piclamilast (picla, 1 µM) brought the ratio to a higher level. The ratio was not further raised by the non-specific phosphodiesterase inhibitor IBMX (200 µM), indicating the saturation level of the biosensor by cAMP (Rmax). These experiments were repeated with similar results (Figure 2D, left; fsk: 0.19; fsk+cilo: 0.49; fsk+cilo+picla: 1.00, expressed as a fraction of Rmax; N=11; n=18; A=7; Friedman’s rank test: F=22, P<0.001; #1 vs #2: P<0.001; #2 vs #3: P<0.001). PDE3 thus contributes importantly to the regulation of cAMP level, and when this phosphodiesterase is inactivated pharmacologically, PDE4 remains to control cAMP level.

We then wondered whether this situation was similar when PDE4 is inhibited first while PDE3 is active. In a similar protocol, on top of forskolin stimulation, PDE4 was first inhibited with piclamilast, which induced an increase in cAMP level in ChINs (Figure 2B). PDE4 activity thus also contributes to the regulation of cAMP level reached upon forskolin application, even though PDE3 is present and active. Indeed, further application of cilostamide increased the ratio to Rmax, the level that was not raised by IBMX. This protocol was repeated with similar results (Figure 2D, middle; fsk: 0.19; fsk+picla: 0.46; fsk+picla+cilo: 1.00; N=6; n=7; A=4; Friedman’s rank test: F=12, P<0.001; #1 vs #2: P=0.016; #2 vs #3: P=0.016).

We explored the contribution of other phosphodiesterases that are expressed in the striatum. Since PDE2A regulates elevated cAMP levels, its activity was tested when adenylyl cyclases were activated and PDE3 was inhibited (Figure 2C). In the presence of forskolin and cilostamide, the ratio level in the presence of the selective PDE2A inhibitor PF-05180999 (PF-05, 1 µM) was slightly increased compared to the previous the steady-state level (Figure 2D; fsk: 0.24; fsk+cilo: 0.44; fsk+cilo+PF-05: 0.48; N=7; n=13; A=6; Friedman’s rank test: F=11.1, P<0.001; Wilcoxon #1 vs #2: P=0.008; #2 vs #3: P=0.046). The amplitude of the response to PF-05 was however considerably smaller than the effect of picla (Wilcoxon-Mann-Whitney two-sample rank test, P=5.10^−5^). PDE10A has a high affinity for cAMP and we wanted to test its potential effect on lower cAMP levels, i.e. in a condition in which PDE3 and PDE4 are active. The PDE10A inhibitor TP-10 (100 nM) had no effect on the forskolin-induced steady-state cAMP level (Figure 2D: N=5; n=7; A=4; Wilcoxon P=0.16), indicative of no PDE10A activity in ChINs.

**Figure 2:**
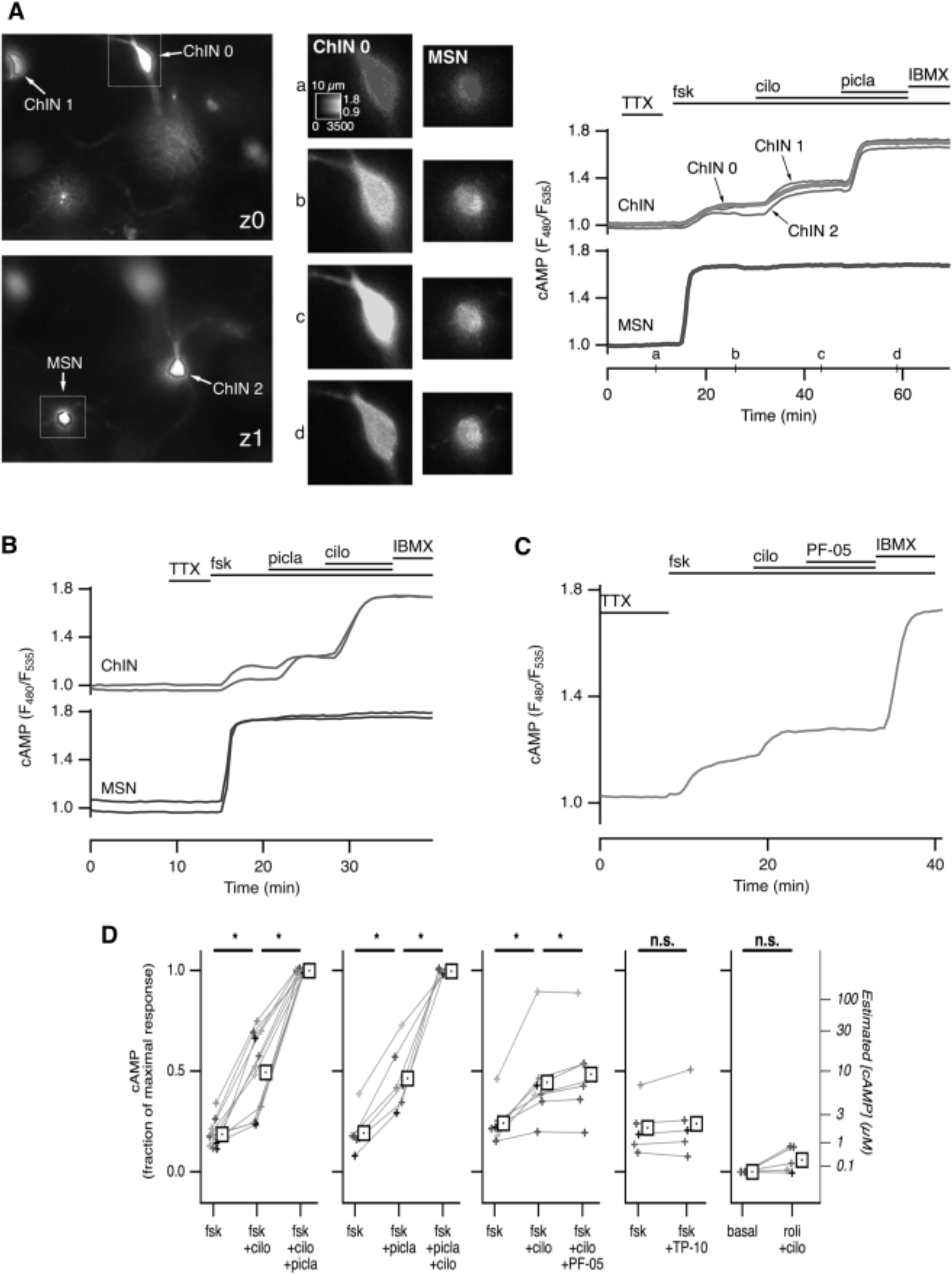
PDE3 and PDE4 are the main phosphodiesterases controlling cAMP in ChINs. (A) The Epac-S^H150^ biosensor for cAMP was expressed in mouse pup striatal brain slices and imaged with wide-field microscopy. Two focal planes (z0 and z1) in the same field of view were acquired at each time-point (a, b, c, d). Regions of interest shown on the grayscale images define the area used to measure the ratio (F480/F535) for 3 individual ChINs (red ROIs, ChIN 0 to 2, and red traces in the graph) and an MSN (blue ROI and blue trace in the graph). Dashed squares indicate the regions shown in pseudocolor for each time-point (a-d) indicated on the graph, and show a ChIN and an MSN. Horizontal bars on the graph indicate the bath application of drugs: tetrodotoxin (TTX, 0.2 µM), forskolin (fsk, 13 µM), cilostamide (cilo, 10 µM), piclamilast (picla, 1 µM) and IBMX (200 µM). (B) Same protocol as in (A) except that the order of application of cilostamide and piclamilast was inverted. (C) Same protocol as in (A) except that, instead of picla, PF-05180999 (PF-05, 1 µM) was applied to inhibit PDE2A. (D) For each experiment, the average normalized ratio is measured over the steady-state ratio for each drug combination. If several ChINs were present in the same recording, their measurements are averaged. For each experiment, the successive measurements are plotted as markers with a same color and connected by a line. The average of the repeated experiments is indicated with a square marker. * indicates a statistically significant difference.

We then wanted to determine whether ChINs displayed a tonic cAMP production. In the absence of forskolin, the combination of rolipram and cilostamide induced no significant ratio change in ChINs (Figure 2D: N=5; n=6; A=3; Wilcoxon P=0.063), indicating that basal cAMP production in these conditions is negligible. This comes in stark contrast with the tonic cAMP production revealed in MSNs when inhibiting PDE10A (46).

Collectively, these experiments show that in ChINs, when cAMP production is stimulated, cAMP levels remain low by the combined action of PDE3 and PDE4.

### The cGMP pathway increases cAMP levels through PDE3 inhibition

PDE3 degrades both cAMP and cGMP, but the Vmax of PDE3 for cGMP is very low and cGMP thus behaves in vitro as a competitive inhibitor of PDE3 activity towards cAMP hydrolysis (26). Furthermore, a PKG-dependent phosphorylation site in PDE3A that negatively regulates PDE3A activity has been reported (47). We therefore pondered whether a cGMP signal, by inhibiting PDE3, could increase cAMP level. We used the NO donor compound DEANO (2-(N,N-diethylamino)-diazenolate 2-oxide) which activates the soluble guanylyl cyclase and increases cGMP level: after forskolin increased cAMP to a steady-state level, the application of DEANO (10 µM) produced a marked cAMP increase. This cAMP level was not further increased by the application of the PDE3 inhibitor cilostamide (Figure 3A and Figure 3B, left panel; fsk: 0.32 of maximal response; fsk+DEANO: 0.71; fsk+DEANO+cilo: 0.73; N=6, n=10, A=5; Friedman’s rank test: F=10.3, P=0.002; Wilcoxon #1 vs #2: P=0.016; #2 vs #3: P=0.22). DEANO thus reproduces and occludes the effect of cilostamide. Conversely, when added on top of forskolin and cilostamide, DEANO induced no further cAMP increase, indicating that cilostamide also occludes DEANO effect (Figure 3B, right panel; fsk: 0.33; fsk+cilo: 0.62; fsk+cilo+DEANO: 0.63, N = 5, n=8, A=4; Friedman’s rank test: F=7.6, P=0.02; Wilcoxon #1 vs #2: P=0.03; #2 vs #3: P=0.6). These experiments demonstrate that the activation of the cGMP pathway in ChINs increases cAMP via PDE3 inhibition.

**Figure 3:**
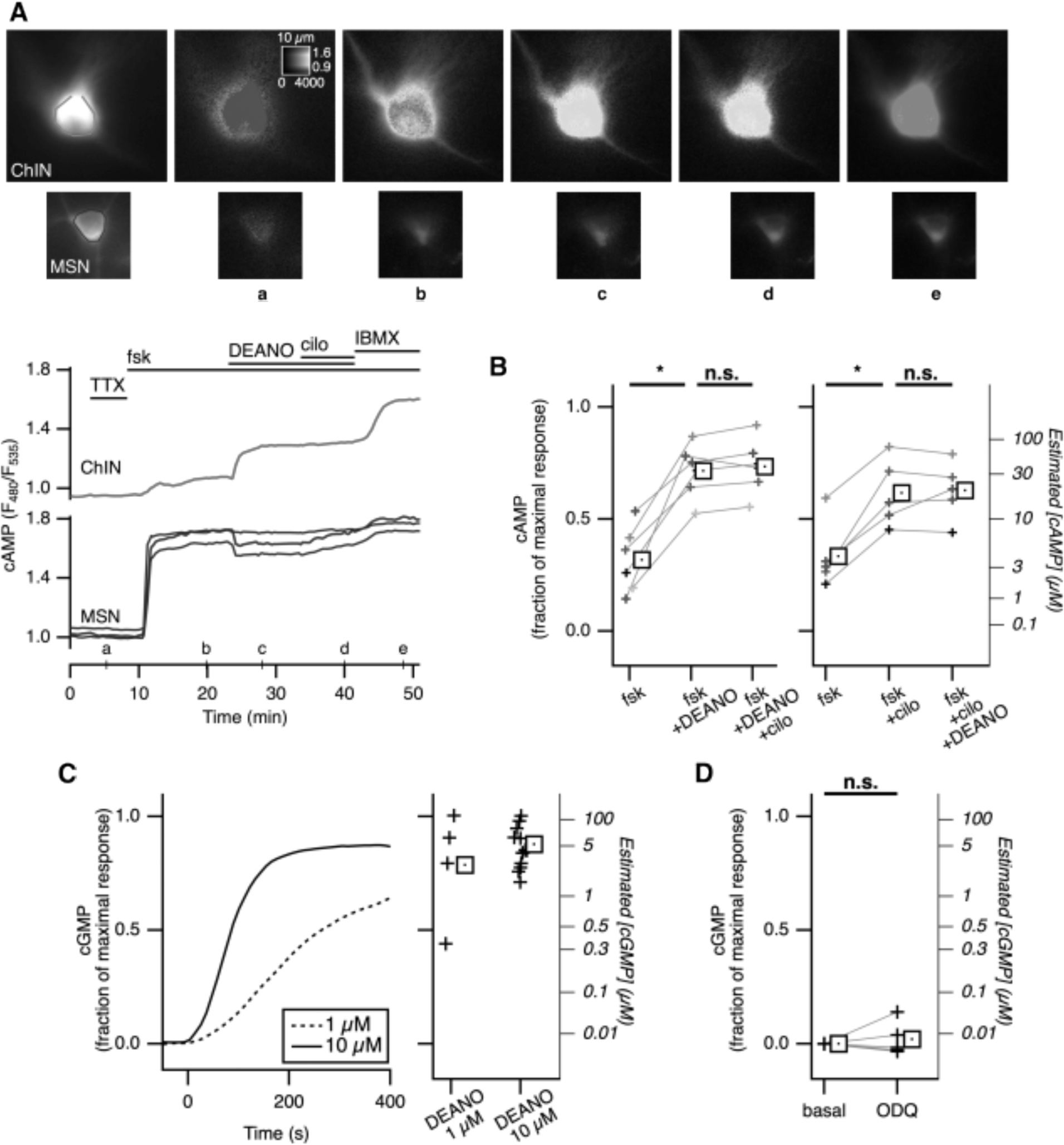
NO donor increases intracellular cAMP through PDE3A inhibition. (A) The Epac-S^H150^ biosensor for cAMP was expressed in striatal brain slices and imaged with wide-field microscopy. Images show a ChIN (top line) and an MSN (bottom line). Traces below show the ratio of these ROIs as well as other MSNs (not shown). (B, left) The same experiment was repeated 6 times. Plots display the average of the response to forskolin 13 µM (fsk) alone, with DEANO 10µM (fsk+DEANO) and cilostamide 10 µM (fsk+DEANO+cilo). (B, right) Average ratio responses when cilo was applied before DEANO. The average of the different repeats is represented with a square symbol. * indicates a statistically significant difference. n.s. = non significant. (C) the NO donor DEANO increases cGMP in ChINs. The cyGNAL biosensor for cGMP was used to monitor changes in intracellular cGMP concentration during the bath application of 1 or 10 µM DEANO. Traces on the left show the average time-course of the ratio increase for 1 or 10 µM DEANO in ChIN. The plot shows the steady-state ratio for each experiment as a cross marker. The average is indicated with a square marker. (D) the inhibitor of soluble guanylyl cyclase ODQ (20 µM) had no effect.

As a control, we verified with the cGMP-selective biosensor cyGNAL (35) that bath application of DEANO (10 µM) induced a rapid and large increase in cGMP concentration in ChINs (Figure 3C). Bath application of the inhibitor of soluble guanylyl cyclase ODQ (20 µM) had no effect (Figure 3D: N=5; n=7; A=1; Wilcoxon P=0.7), showing a lack of basal cGMP production in our experimental conditions.

Altogether these findings unequivocally establish that cGMP signaling increases cAMP levels in ChINs by suppressing PDE3 activity. This signaling mechanism strikingly contrasts with cGMP effects in MSNs where it activates PDE2A and thus reduces cAMP level (36).

### The calcium pathway decreases cAMP and cGMP via PDE1 activation

We previously observed that, in MSNs, PDE1 activity efficiently regulates cAMP and cGMP signals in the presence of calcium (35) illustrating that PDE1 is an integrator of cyclic nucleotide and calcium signaling. This effect in MSNs is likely mediated by the PDE1B isoform (48). PDE1A isoform is expressed in ChINs (28) but its functional role has remained unexplored. Here, we tested whether PDE1A activity in ChINs might down-regulate cAMP and/or cGMP signals when activated by calcium. First, we established reliable protocols to increase intracellular calcium. We used the Twitch-2B ratiometric biosensor to measure the calcium signal triggered by NMDA receptors and type I metabotropic glutamate receptors (mGlu1/5) (Figure 4). NMDA receptors were activated transiently by NMDA released from the “caged” precursor MNI-NMDA (100 µM) by a flash of UV light, as previously described (35). This protocol triggered a large and transient calcium increase in all ChINs tested, as well as in MSNs if present in the same field of view (Figure 4A). Figure 4B shows the calcium response to NMDA in ChINs with the average trace (left), and baseline and peak amplitude (right) for individual ChINs. We also observed that mGlu1/5 receptor activation by bath application of the selective agonist DHPG (50 µM), triggered a calcium increase of large amplitude and duration (Figure 4A, C). Interestingly, while mGlu1/5 receptors are also expressed in MSNs, no or little calcium response to DHPG was observed in these neurons.

**Figure 4:**
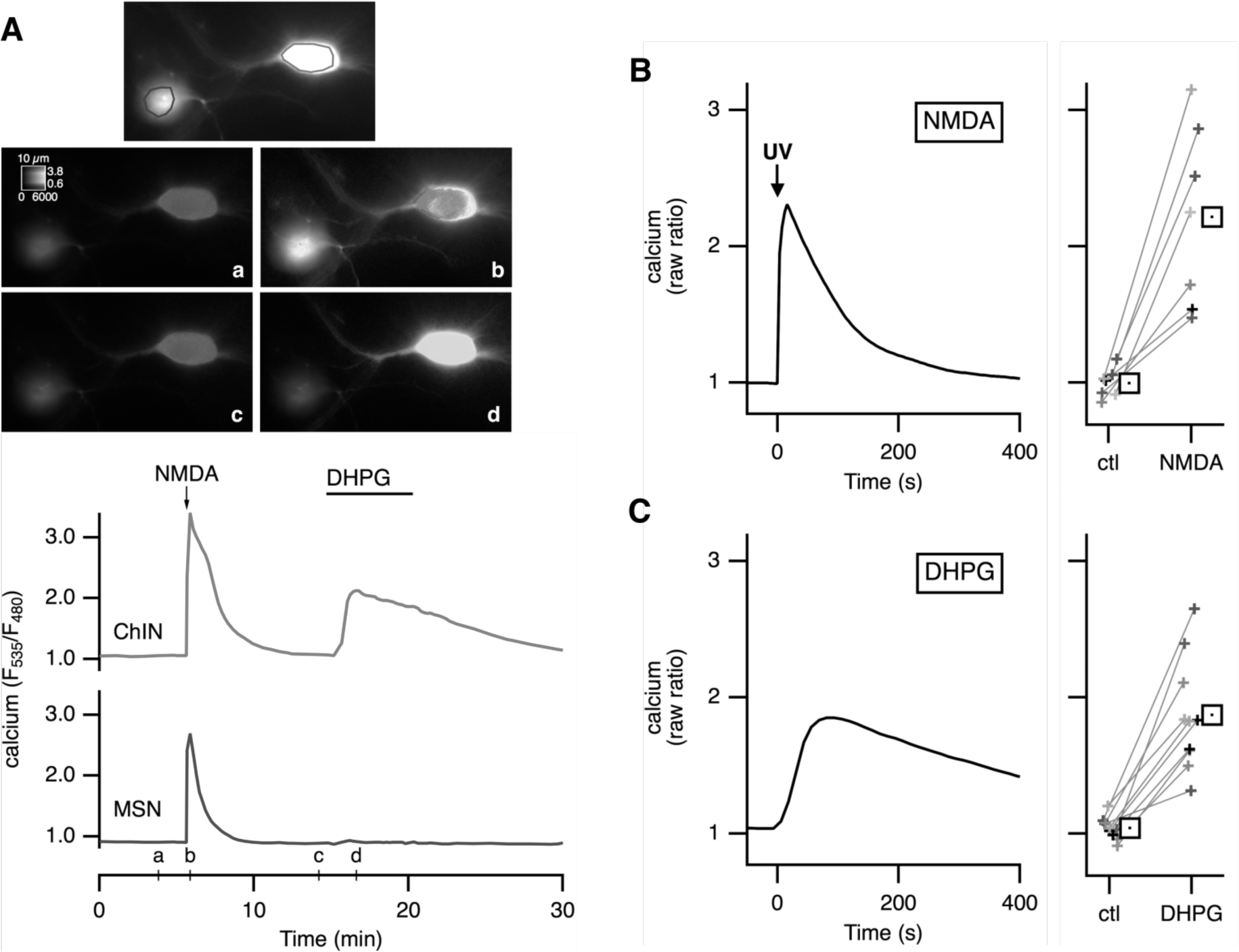
NMDA photorelease or activation of group I mGlu receptors induces large calcium signals in ChIN. The Twitch-2B calcium biosensor was expressed in striatal brain slices and imaged with wide-field microscopy. (A) Intracellular calcium is monitored over regions of interest drawn on a ChIN and on an MSN (red and blue, respectively). Release of NMDA from MNI-NMDA with a flash of UV light (arrow) raised calcium in both neurons, whereas sustained bath application of 50 µM DHPG, an agonist of group I metabotropic glutamate receptors, increased intracellular calcium levels selectively in ChIN. (B, C) The average time-course of the ratio change triggered by NMDA and DHPG in ChIN is shown on the left; plots on the right display the baseline and peak ratio value measured in ChINs for independent experiments (B: N=4 ; C: N=5). The average of the different repeats is represented with a square symbol.

We then tested whether such calcium increases activate PDE1 and therefore could reduce cAMP or cGMP. A steady-state elevated cAMP level was first obtained using forskolin and blocking PDE3 and PDE4 (with either 1 µM cilo and 1 µM roflumilast or 10 µM cilo and 1 µM piclamilast) and PDE2A (with PF-05180999, 1 µM). In order to stimulate a moderate cAMP production and thus maintain the visibility of PDE1 action, a lower concentration of forskolin (0.5 µM) was employed in these experiments. NMDA uncaging or DHPG bath application both produced large decreases in cAMP levels in ChIN (Figure 5 A,C,E: before NMDA application : 0.88 of Rmax, NMDA application : 0.29, N=6, n=9, A=5; before DHPG: 0.77, DHPG application: 0.40, N=5, n=7, A=4). These effects were largely reduced with Lu AF64196 (1-10 µM), a potent and selective PDE1 inhibitor (49) for NMDA (before NMDA application : 0.91 of Rmax, NMDA application : 0.83, N=6, n=9, A=4; ratio change significantly reduced by Lu AF64196, Wilcoxon rank test, *P*=8.10^−4^) and for DHPG (before DHPG: 0.86, DHPG application: 0.89, N=5, n=8, A=3; ratio change significantly reduced by Lu AF64196, Wilcoxon rank test, with *P*=0.002). It should also be noted that, in the presence of the PDE1 inhibitor Lu AF64196, some changes in cAMP level still remained, which can result either from incomplete PDE1A inhibition and/or from NMDA effects on other targets such as calcium-modulated adenylyl cyclases.

**Figure 5:**
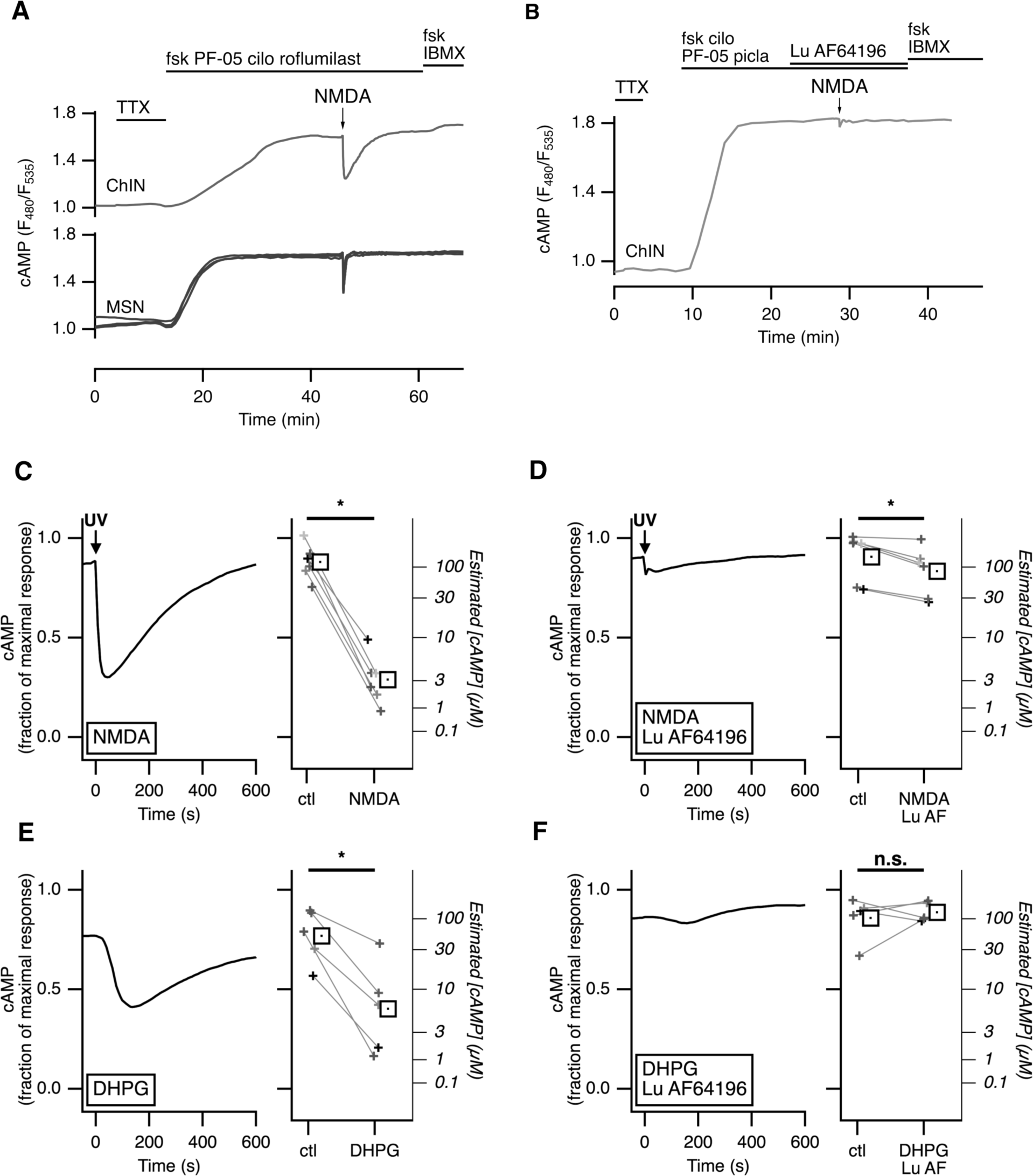
activation of NMDA or mGlu1/5 receptors in ChIN decreases cAMP via PDE1 activation. The Epac-S^H150^ biosensor for cAMP was expressed in striatal brain slices and ChINs were imaged with wide-field microscopy. (A) cAMP production was stimulated with 0.5 µM fsk in the presence of inhibitors of PDE2A (PF-05180999, 1 µM), PDE3 (cilo, 1 µM) and PDE4 (roflumilast, 1 µM). On the steady-state cAMP level, a flash of UV light at the time indicated by the arrow photoreleased NMDA from MNI-NMDA (100 µM), triggering a ratio decrease, in both ChIN and MSNs. (B) The selective PDE1 inhibitor Lu AF64196 (1 µM) prevented this decrease. C-F: these experiments were repeated using phosphodiesterase inhibitors (PDE3: cilo, 1 µM or 10 µM; PDE4: roflumilast or piclamilast, 1 µM), without (C) or with Lu AF64196 (1 or 10 µM) (D). The graph shows the average normalized cAMP signal in ChINs, and the plot shows the amplitude of the steady-state cAMP level (ctl) and minimal cAMP level reached in response to NMDA photorelease (NMDA). A similar set of experiments was performed except that DHPG (50 µM) was applied in the bath. This also induced a transient decrease in cAMP concentration (E). This decrease was suppressed by Lu AF64196 (F).

We then tested whether the activation of PDE1 by calcium could also regulate cGMP level in ChINs. In these experiments, NMDA receptors were transiently activated by the MNI-NMDA uncaging protocol, and the mGlu1/5 receptors were activated with bath application of 1 µM quisqualate (AMPA and mGlu receptors agonist) in the presence of the AMPA receptor antagonist NBQX (1 µM) and NMDA antagonist D-AP5 (20 µM). First, a steady-state cGMP level was obtained with a bath application of 10 µM DEANO. NMDA uncaging or activation of mGlu1/5 receptors both produced large decreases in cGMP levels in ChIN (Figure 6, left: before NMDA application : 0.80 of Rmax, NMDA application : 0.18, N=9, n=13, A=9; before quisqualate: 0.83, quisqualate application: 0.22, N=5, n=9, A=4). Again, these effects were largely reduced with Lu AF64196 (1-10 µM) for NMDA (figure 6, right: before NMDA application : 0.76 of Rmax, NMDA application : 0.80, N=6, n=7, A=6; ratio change significantly reduced by Lu AF64196, Wilcoxon rank test *P*=2.10^−4^) and for quisqualate (before quisqualate: 0.88, quisqualate application: 0.84, N=5, n=7, A=3; ratio change significantly reduced by Lu AF64196, Wilcoxon rank test, with *P*=0.002).

**Figure 6:**
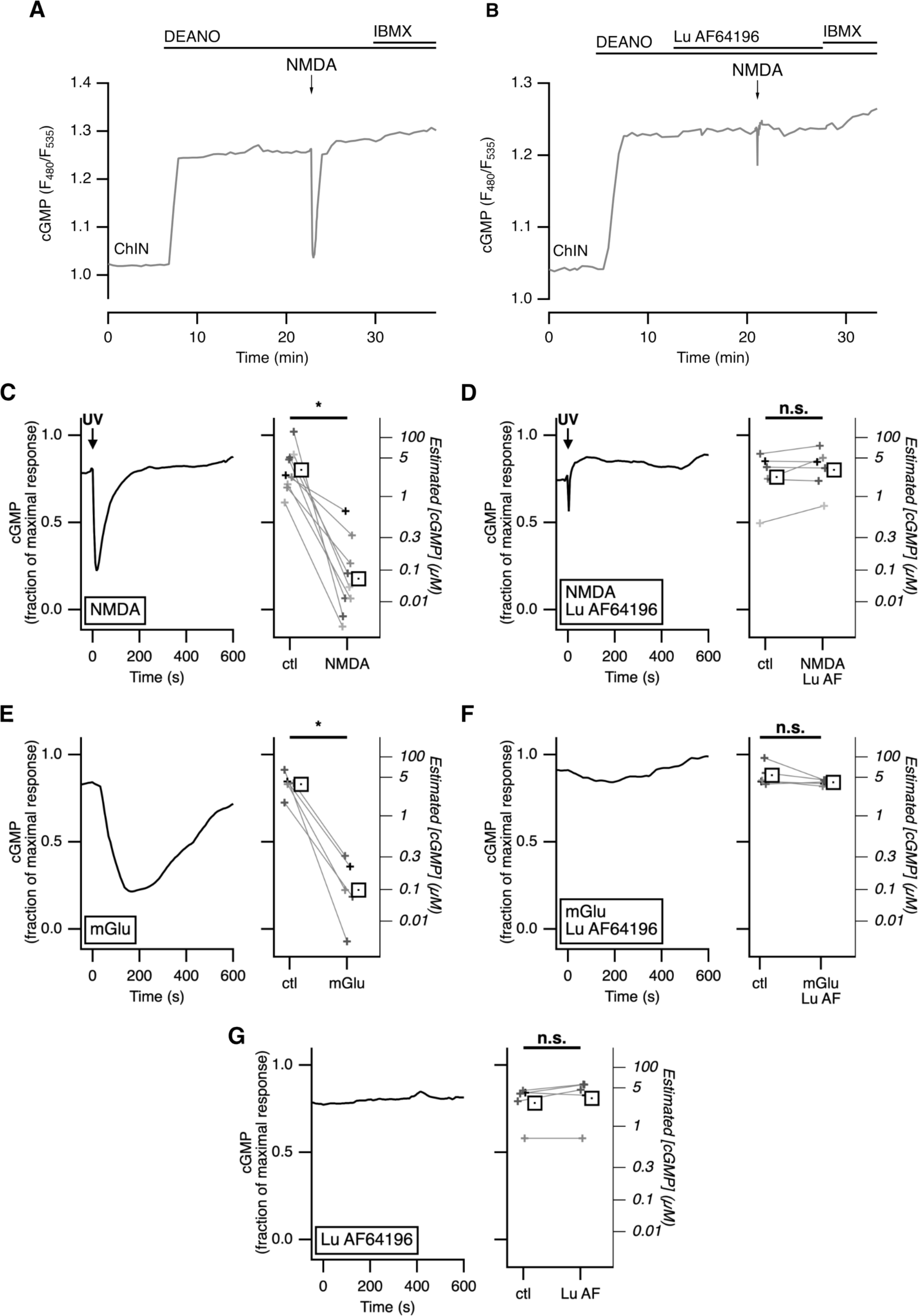
activation of NMDA or mGlu1/5 receptors in ChIN decreases cGMP via PDE1 activation. The cyGNAL biosensor for cGMP was expressed in striatal brain slices and ChINs were imaged with wide-field microscopy. (A) cGMP production was stimulated with 10 µM DEANO. On the steady-state cGMP level, a flash of UV light at the time indicated by the arrow photoreleased NMDA from MNI-NMDA (100 µM), triggering a ratio decrease. (B) The selective PDE1 inhibitor Lu AF64196 (10 µM) prevented this decrease. These experiments were repeated without (C) or with Lu AF64196 (1 or 10 µM) (D). The graph shows the average normalized cGMP signal, and the plot shows the amplitude of the steady-state cGMP level (ctl) and minimal cGMP level reached in response to NMDA photorelease (NMDA). A similar set of experiments was performed except that mGlu1/5 receptors were activated by bath application of quisqualate (1 µM) together with AMPA receptor antagonist NBQX (1 µM) and NMDA antagonist D-AP5 (20 µM). This also induced a transient decrease in cGMPconcentration (E). This decrease was also suppressed by Lu AF64196 (F). (G) lack of effect of Lu AF64196.

As a control, we verified that PDE1 inhibition had no effect on the steady-state cGMP level elicited by 10 µM DEANO (Figure 7G: in DEANO: 0.78 of Rmax; in DEANO and Lu AF64196: 0.81; N=5, n=6, a=5; Wilcoxon P=0.094).

**Figure 7:**
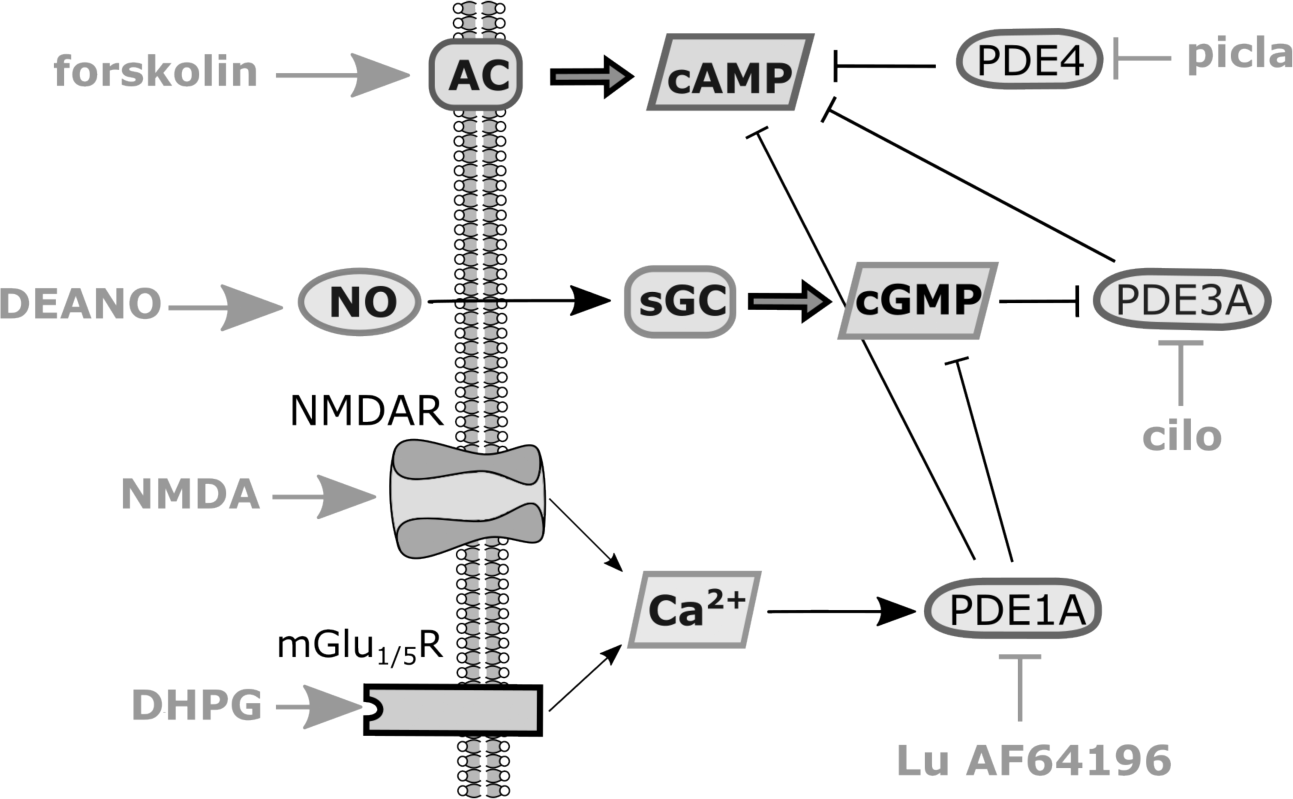
schematic representation of phosphodiesterases and cross-pathway regulations in ChINs.

Collectively, these experiments show that ChINs express functional PDE1 that efficiently degrades both cAMP and cGMP in response to an increase in intracellular calcium.

## Discussion

### ChINs *vs*. MSNs integrative properties

Our study highlights striking differences between ChINs and MSNs in the way they handle cAMP signals: while MSNs reach a rapid and maximal response with forskolin, ChINs showed slow and moderate increases in cAMP concentration. These observations reinforce the view that ChINs and MSNs bear intrinsic differences in signal integration properties that endow them with fundamentally different functions in the striatal network. Among these differences, our experiments highlighted how the NO-cGMP pathway is differentially integrated in ChINs and in MSNs through the expression of different phosphodiesterases.

### Phosphodiesterases in ChINs

Phosphodiesterases determine the shape and dynamics of cAMP and therefore the specificity of cAMP signal integration. Previous transcriptomic and proteomic studies suggested the expression of specific phosphodiesterases in striatal ChIN (21, 22, 28) but no functional analysis has been performed so far. We used highly specific phosphodiesterase inhibitors to acutely test the functional contribution of these phosphodiesterases. Our experiments demonstrate for the first time that PDE1, PDE3 and PDE4 are functional in striatal ChINs, and regulate the dynamics of cAMP in the context of NO or glutamate neuromodulation (Figure 7). This specific combination of functional phosphodiesterases contrasts with the situation in MSNs which express PDE1B, PDE2A, PDE4 and PDE10A. In our experiments, we observed little or no signs of PDE2A or PDE10A activity in ChINs, consistent with the documented absence of their mRNA in these cells (22, 33, 34). This stands in contrast with the important functional role played by PDE2A and PDE10A in MSNs (36, 46, 50).

Altogether, PDE3A emerges as a characteristic feature of ChINs, along with the expression of PDE1A and PDE4.

### Cross-pathway integration: cAMP and cGMP

Remarkably, since PDE3 is inhibited by cGMP, its functional presence in ChINs suggests potential cross-regulation of cAMP by the NO-cGMP signaling pathway, which is demonstrated by our experiments. Surprisingly, this interesting possibility had never been explored in neurons, while its demonstration in other physiological conditions remains scarce (23, 51, 52). This mechanism has also been described in cardiomyocytes, with the added complexity of PDE2A being co-expressed with PDE3: depending on cAMP levels, NO donors either potentiate or reduce I_Ca_, consistent with the inhibition of PDE3 or the activation of PDE2A. This effect has been reported in frog ventricular cardiomyocytes (53) as well as in human cardiomyocytes (54, 55).

Pharmacological inhibition of PDE3 is used in the treatment of some clinical conditions of the cardiovascular system. PDE3 inhibitors, such as milrinone, exert powerful positive inotropic and lusitropic effects and are used in the treatment of acute heart failure (56). Milrinone has also been shown to prevent delayed ischemic injury in the brain (57–59). However, while several preclinical and clinical studies showed beneficial effects of PDE3 inhibitors in neuronal functions, whether these effects result from a direct action on neurons or indirect effects through the improvement of vascular condition remains uncertain.

Our data show for the first time in neurons that the effect of NO-cGMP neuromodulatory axis positively controls cAMP levels via PDE3 inhibition. This mechanism may be implicated in the excitatory effect of NO donors documented in ChIN, an effect probably mediated by the modulation of I*_h_* current by cAMP signaling (60–62). This highlights a novel mechanism of cross-pathway integration in the striatum. The NO/cGMP pathway has long been known to play an important role in striatal function by regulating excitability, dopamine release and synaptic transmission. Our work highlights an entirely new mechanism by which NO-induced PDE3 inhibition would mediate an increase in acetylcholine release by ChINs. Importantly, this NO-mediated increase in cAMP levels observed in ChIN sharply contrasts with its effect in MSN, where, by stimulating PDE2A, NO donors down-regulate cAMP signaling (36).

### Cross-pathway integration: cyclic nucleotides and calcium

Another cross-pathway integration level is provided by PDE1, which powerfully degrades both cAMP and cGMP when activated by calcium-calmodulin (69). Calcium imaging experiments showed that while activating NMDA receptors increased intracellular calcium, ChINs also displayed a powerful and sustained calcium signal in response to the activation of group I metabotropic glutamate receptors. This is consistent with the expression of mGlu1 and mGlu5 receptors (70–73), which are known to couple through G_q_/G_11_ to the release of intracellular calcium stores. In this study, the stimulation of both NMDA and group I metabotropic receptors efficiently reduced intracellular cAMP and cGMP, an effect similar to what was reported previously in MSNs (35). We hypothesize that this effect is mediated by PDE1A in ChINs, whereas PDE1B is expressed in MSNs (22, 28).

In the last decade, the pharmacological inhibition of PDE1 was of considerable interest, particularly in the context of CNS diseases (74). Among the various PDE1 inhibitors under investigation, Lu AF64386 has shown promising results in reduction of impulsivity and reversal of neurophysiological deficits in the rat phencyclidine model of schizophrenia (75), indicating that PDE1 inhibition may be beneficial in disorders implicating a dysfunction of the medial prefrontal cortex and nucleus accumbens network. Another line of research has highlighted PDE1 inhibitors as pro-cognitive and memory enhancing (76, 77). PDE1 inhibitors have also demonstrated anti-parkinsonian effects, particularly potentiating Levodopa effect in rodent and monkey models of Parkinson’s disease, as well as having potent efficacy against Levodopa induced dyskinesia (78). In a phase 1b research on PD in humans, a PDE1 inhibitor has shown therapeutic effects, and currently, Lenrispodun is in Phase 2 clinical trial for the treatment of Parkinson’s disease (NCT05766813).

Our data support the notion that PDE1 plays a role in the integration of calcium and cyclic nucleotide signaling in ChINs. In other types of neurons, PDE1 has been shown to control the calcium threshold for synaptic plasticity (79), but the precise physiological function of cyclic nucleotide and calcium signaling in ChINs is still unknown. Further investigations combining multifaceted techniques and behavioral assays are needed to understand the influence of PDE1A and PDE1B isoforms on neuronal networks and behavior.

### Signaling pathways intricacies

The co-expression of PDE1A and PDE3A in ChINs raises the intriguing question of what happens in physiological situations when glutamatergic inputs occur simultaneously with NO release: activation of PDE1A by calcium is expected to reduce cAMP levels, while PDE3A inhibition by cGMP should increase cAMP. PDE3A has a sub-micromolar Km for cAMP (80) whereas PDE1 has a much higher Km value (81), suggesting that, on low cAMP levels, cGMP-mediated inhibition of PDE3A should win against PDE1A and therefore raise cAMP level. However, PDE1A also degrades cGMP, which implies that it may temper the impact of NO on the cAMP pathway. Predicting the precise interplay of these effects within ChINs thus appears challenging. Furthermore, the possibility of phosphodiesterases residing in distinct sub-cellular signaling domains can substantially alter their overall impact. Ultimately, the integrated response will be shaped by the kinetics governing these events relative to one another.

The low responsiveness of ChINs to cAMP-activating signals such as forskolin is striking but the underlying cellular mechanisms remain to be determined. All adenylyl cyclases except AC9 are activated by forskolin (82). AC1, AC2 and AC5 are widely expressed in the brain, including the striatum (83). Cluster analysis of mRNA transcript suggest that AC1 and AC2 predominate in ChINs whereas AC5 would dominate in MSNs (22). This indicates that the reduced responsiveness to forskolin in ChINs compared to MSNs does not result from ChINs lacking adenylyl cyclases sensitive to forskolin. Another important factor is the amount of active signaling enzyme, i.e. the density of cyclase per membrane surface or amount of calcium-store per cytosolic volume, in relation with cell geometry. Indeed, ChINs have a large soma, their proximal dendrites have a large diameter compared to the much thinner MSNs (45, 84). Surface to volume ratio critically determines whether a signal gets amplified or dampened (15) and in the case of cAMP, small neurons like MSNs can accumulate cAMP more easily. In contrast, ChINs with a larger cytosol in which cAMP gets degraded by PDEs are in a less favorable position to accumulate enough cAMP to be detected by our cytosolic biosensors.

The way in which these cellular events come together within the network remains poorly understood. However, our data strongly suggest that this new cGMP-mediated up-regulation of cAMP signaling should be considered in physiological scenarios. Somatostatin- and NPY-positive interneurons also express NO synthase and are the primary NO source in the striatum (45, 84). The release of NO depends on intracellular calcium levels and is heightened by dopamine (66, 85). The convergence of glutamatergic and dopaminergic inputs is thus expected to trigger the release of NO, which should favor ChINs excitability via the cGMP/PDE3A/cAMP mechanism. Whether this mechanism contributes to the rebound excitation observed in ChINs after phasic dopamine release, or contributes to the broader regulation of ChIN activity should be a focus of future research.

These observations may be of particular significance in the context of pathological conditions such as Parkinson’s disease. Indeed, the first medications for Parkinson’s disease were based on anticholinergic drugs, and there is compelling evidence that photoinhibition of ChIN has prominent anti-parkinsonian effects (86, 87). A noteworthy observation is that the NO-mediated increase in ChIN firing is markedly more pronounced in a mouse model of Parkinson’s disease (61). This suggests that moderating the excessive ChIN activity might reduce some symptoms of Parkinson’s disease. This novel understanding of the intricate interplay between NO/cGMP, PDE3A and cAMP opens interesting possibilities for the development of more effective and precisely tailored therapeutic strategies in the future.

## Abbreviations

ChIN: cholinergic interneuron
MSN: medium-sized spiny neuron
NO: nitric oxide

## Acknowledgements

We would like to gratefully thank Drs Corinne Beurrier, Julie Perroy, Jérémie Naudé and Pr Juan Llopis for their careful reading of this manuscript and helpful comments.

